# Can the response of above- and below-ground traits to overgrazing vary between annual and perennial species?

**DOI:** 10.1101/771444

**Authors:** Hui Zhang, Wenxing Long

**Author notes:** **Correspondence author:** Hui Zhang.

## Abstract

Understanding functional traits that capture species response to grazing is crucial for uncovering mechanisms underlying the grazing-induced alterations in vegetation composition. Currently, the majority of studies ignore the effects of life form on the response of above- and below-ground traits to grazing. In this study we quantified i) whether below-ground traits exhibit similar response to overgrazing as do above-ground traits; and ii) whether the response of above- and below-ground traits to overgrazing varied with life form (annual versus perennial species). To testify the generality of these two questions, we compared common grassland species between an undisturbed and an over grazed meadow at each of four elevations ranging from 3000m to 3600m on the Qinghai Tibetan Plateau. We found both above-ground and below-ground traits exhibited similar resource conservation *vs*. resource acquisition trade-off for responding to overgrazing. Moreover, perennial species but not annual species determine the response of above- and below-ground traits to overgrazing. This work bridges the gap in the literature for the influence of land-use (in the absence and presence of overgrazing) and life forms (annual and perennial species) on species-specific response at both above-, and below-ground level.

## Introduction

Grazing is one among the most significant drivers of global vegetation composition (Díaz *et al*., 2007; Stein *et al*., 2016). Understanding functional traits that capture species response to grazing is crucial for uncovering mechanisms underlying the grazing-induced alterations in vegetation composition (Díaz *et al*., 2001; Cingolani *et al*., 2005; de Bello *et al*., 2005; Díaz *et al*., 2007; Evu *et al*., 2009; Golodets *et al*., 2010). Generally, above-ground traits such as shorter plant height and higher specific leaf area (SLA) can help the plants have high resource acquisition for tolerating high grazing pressure (Díaz *et al*., 2001; Li *et al*., 2017). In contrast, in the undisturbed condition, species prefer to have lower SLA but higher plant heights, which is regarded as a strategy for high resource conservation to adapt to resource limitation (Westoby, 1999).

It has also been found that below-ground traits play a key role in mediating grazing-induced alterations to vegetation composition (Klumpp *et al*., 2009; McInenly *et al*., 2010). However, due to the notable lack of relevant below-ground trait data (Bergmann *et al*., 2017), it remains unclear whether below-ground traits exhibit similar high resource acquisition (e.g., higher RD, SRL, SRA, but lower RTD) for tolerating grazing as do above-ground traits (Craine *et al*., 2001; Hodge, 2004; Klumpp *et al*., 2009; McInenly *et al*., 2010; Roumet *et al*., 2016; Kramer *et al*., 2016; Feng *et al*., 2018). This limits our understanding of how livestock influence vegetation compositions (Li *et al*., 2017).

Determining the response of above-ground and below-ground traits against grazing requires an understanding of how grazing influences species composition (Díaz *et al*., 2001; James *et al*., 2001). A global meta-analysis showed that overgrazing promotes annual species over perennial species (Díaz *et al*., 2007). Conversely, without grazing pressure, perennial species are expected to dominate (Zhang *et al*., 2018). This is likely due to the fact that annual species tend to have above- and below-ground traits that are associated with high resource acquisition, whereas perennial species tend to possess traits related to a resource conservation strategy (Roumet *et al*., 2006, 2016; Zhang *et al*., 2018). As a result, the response of above-ground and below-ground traits to overgrazing may vary between annual and perennial species. However, the effect of life form (annual versus perennial species) on overgrazing induced alterations to above- and below-ground traits have been hardly considered.

Here, our goal was to test i) whether above-, and below-ground traits have similar response to overgrazing; and ii) whether the response of above-, and below-ground traits to overgrazing differs between annual and perennial species. To tesify the generality of these two goals, we compared above-, and below-ground traits for all common species found in one undisturbed meadow and one overgrazed meadow at each of four elevations ranging from 3000 to 3600 m in Qinghai Tibetan plateau. We hypothesized that 1) both above-, and below-ground traits exhibited a resource conservation *vs*. resource acquisition trade-off for responding to overgrazing; and 2) the response of above- and below-ground traits to overgrazing varies between annual and perennial species.

## Methods and materials

### Study site

The maximum carrying capacity for cold season is approximately 2 yaks per hectare on Qinghai-Tibetan Plateau (Dong *et al*., 2015). In light of this standard, four overgrazed meadows that have undergone overgrazing at a comparable intensity (e.g., overgrazed by approximately the same number of yaks (approximately 4 yaks per hectare) in cold season every year) for nearly 30 years along an increasing elevation ranging from 3000 to 3600 m above mean sea level in the eastern part of the Qinghai-Tibetan Plateau, distributed in Hezuo city, Gansu province, China (34°55′N, 102°53′E; 3000 a.s.l.), Ruergai County, Sichuan province, China (34°1′N, 102°43′E; 3200a.s.l.) and Maqu County, Gansu province, China (34°5’N, 102°10′E; 3400 and 3600a.s.l.). At each elevation, we also sampled one undisturbed meadow for comparison. All sampling regions are characterized by a cold and dry climate; with a mean annual temperature of 2.4 to 3.2 °C. Mean annual precipitation of 530 to 560 mm is primarily distributed from July to August. The local vegetation in undisturbed sites is dominated by herbaceous species, such as *Elymus nutans* (Griseb), *Kobresia pygmaea* (C. B. Clarke), and *Thermopsis lanceolata* (R. Br) (Zhang *et al*., 2015). Soils are classified as alpine meadow soils (Zhang *et al*., 2018).

### Field sampling

Sampling was conducted in August, during the peak of the growing season in 2017. At each of the four elevations, thirty 0.5 × 0.5 m^2^ plots were arranged along four parallel transects in the fenced area and the adjacent overgrazed area of one meadow, with a 20 m spacing between the closest edges of the two areas. The plots were spaced at 20 m intervals. For each of the 30 plots, species compositions was recorded.

### Plant above-ground and root traits measurements

All common species sharing between undisturbed and grazed meadows were sampled to measure three above-ground traits, leaf photosynthesis rate (A;; μmol m^-2^ s^-1^), specific leaf area (SLA; cm^2^/g) and above-ground maximum height (H; m). Further, five root traits, root average diameter (RD; cm), root biomass (RB; g), specific root length (SRL; cm/g), root tissue density (RTD; g/m^-3^), specific root area (SRA; cm^2^/g) were recorded. All of these traits could be good predictors of resource acquisition vs. resource conservation trade-off (Díaz *et al*., 2001; Roumet *et al*., 2006). Traits were measured from three to five mature individuals per species and detail measurement was described in Bergmann *et al*. (2017) and Zhang *et al*. (2018) and Supplementary materials.

## Statistical methods

First, we tested whether there were differences in the above- and below-ground traits for all common species shared between undisturbed and overgrazed meadows at all four elevations using a Wilcoxon signed-rank test. Next, a Spearman correlation test was used to examine the bivariate relationships among all above- and below-ground plant traits at all elevations. Finally, a principal component analysis (PCA) was employed to evaluate which of the eight above-ground and root traits play an important role in affecting species distributions between undisturbed and overgrazed meadows at all elevations. Given the unique set of traits between annual and perennial plant species (Roumet *et al*., 2006; Zhang *et al*., 2018), the above analyses were also performed separately for annuals and perennials to understand whether the response of above- and below-ground traits to overgrazing vary between annual and perennial species.

## Results

In total, we sampled 53-77 species in the undisturbed meadows from 3000 to 3600m elevation, of which 12-22 were annuals and 41-55 were perennial species. On the other hand, only 40-51 species, 22-24 annual and 18-27 perennials, could be identified in the sampling of the overgrazed meadow (Table S1). However, species composition was similar between the undisturbed and overgrazed meadows as the two meadows shared 22-42 species (12-15 annual and 10-27 perennial species) in common (Table S1).

The Wilcoxon signed-rank test showed that all shared species had significantly lower root biomass (RB), root tissue density (RTD), and above-ground maximum height (H) in the overgrazed meadows than those in the undisturbed meadows at all four elevations. In contrast, overgrazed meadows had higher root average diameter (RD), specific root length (SRL), specific root area (SRA), leaf photosynthesis rate (A) and specific leaf area (SLA) than those in the undisturbed meadow (Fig. 1).

**Fig. 1.**
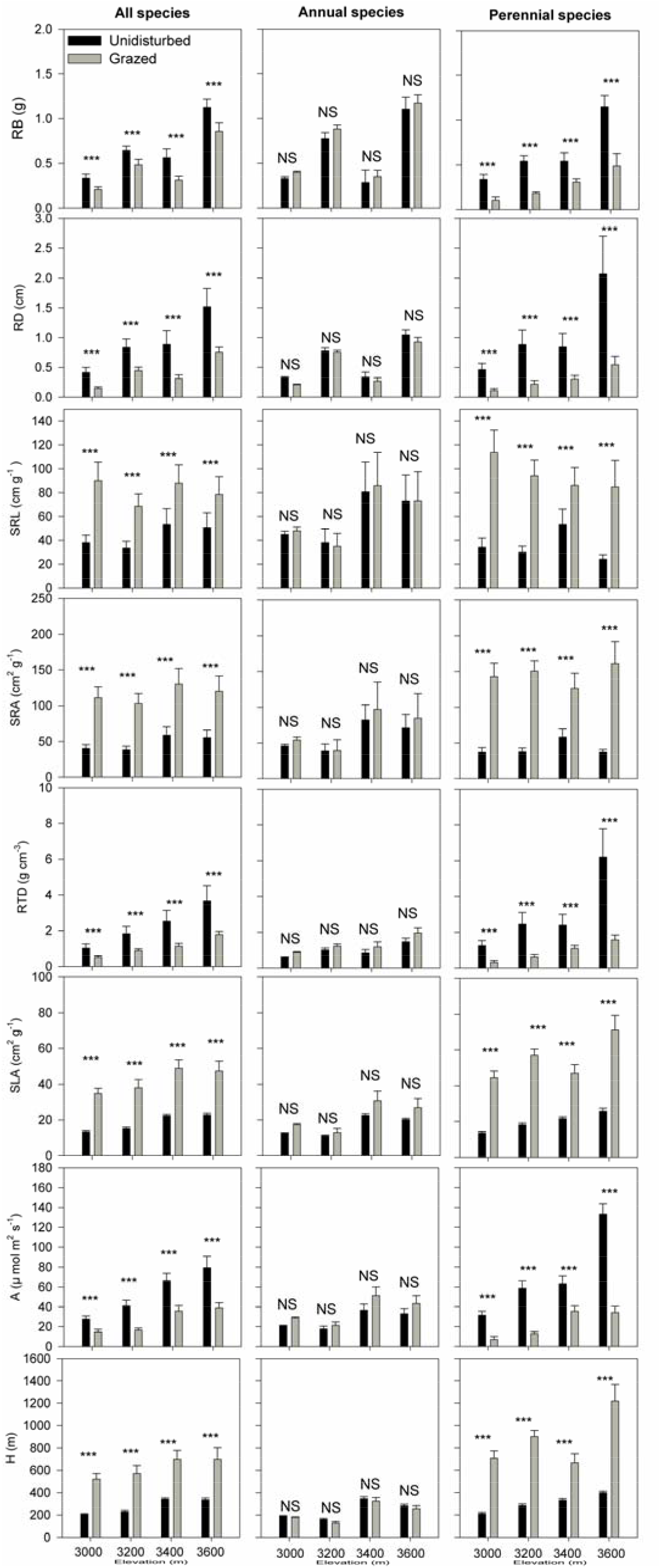
The differences in above-ground traits (height (H; m), leaf maximum −2 −1 2 photosynthesis rate (A; μmol m^-2^ s^-1^) and specific leaf area (SLA; cm /g)) and below-ground traits (root average diameter (RD; mm), root biomass (RB; g), specific root length (SRL; cm^1^/g), specific root area (SRA; cm^2^/g) and root tissue density (RTD; g/m^-3^)) between undisturbed and overgrazed meadows for all species and separately for annual and perennial species from 3000 to 3600m elevations. *** and NS (non-significant) indicates P<0.001 and P>0.05, based on Wilcoxon signed-rank tests.

Consistent results could still be observed in perennial species, when we performed Wilcoxon signed-rank test separately for annuals and perennials (Fig. 1). However, for annual species, there are not any differences in all above-ground and below-ground traits between undisturbed and overgrazed meadows (Fig. 1).

We observed strong correlations amongst the five root traits, as well as between the three above-ground traits, for all common species in both, undisturbed and grazed meadows at all four elevations (Fig. 2). However, when we tested these correlations separately for annuals and perennials, consistent results could still be observed in perennial species in both undisturbed and overgrazed meadows at all elevations (Fig. 2). However, for annual species, most of these strong correlations were non-significant in both undisturbed and overgrazed meadow at all elevations (Fig. 2).

**Fig. 2.**
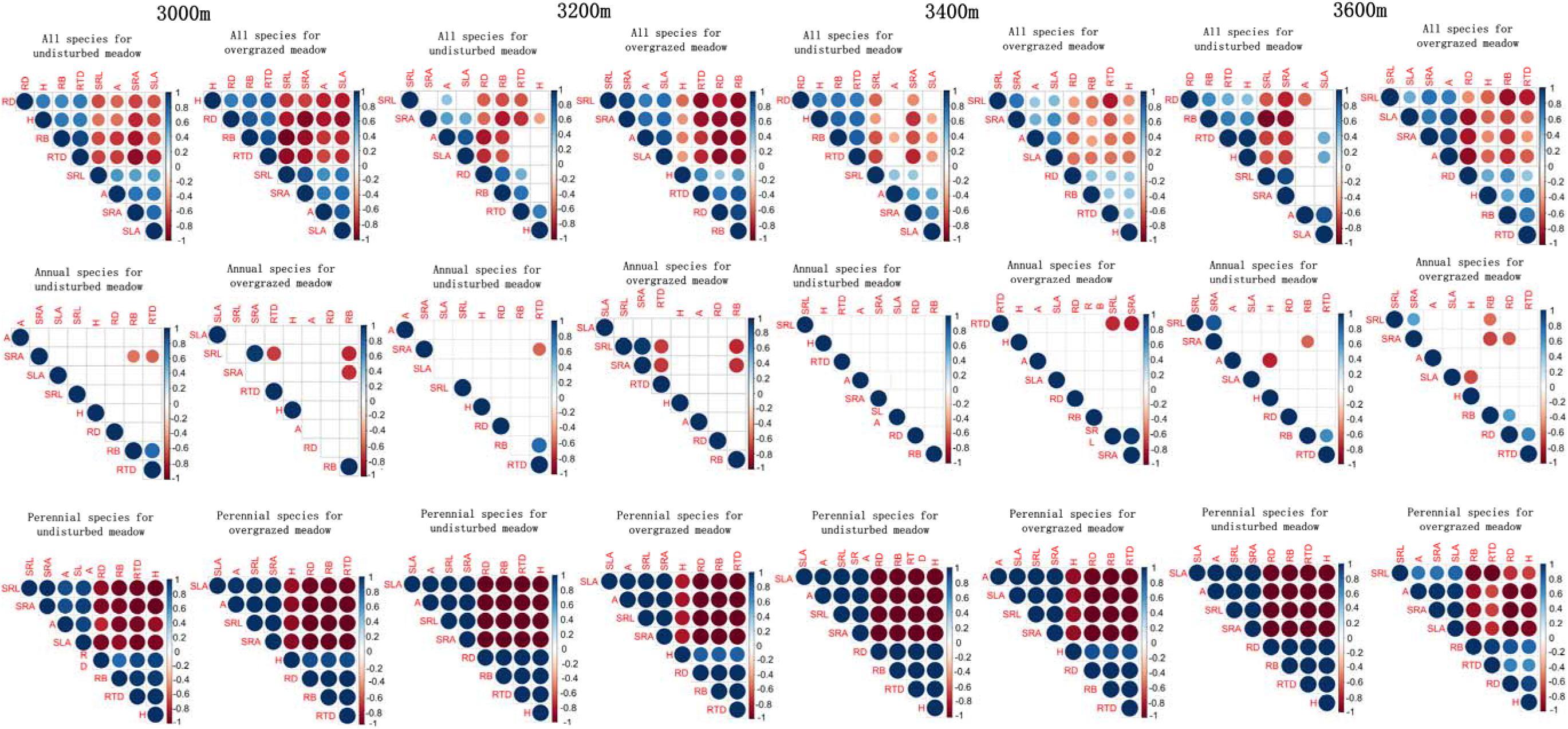
The interrelationships among above-ground traits (lmaximum height (H), leaf maximum photosynthesis rate (A) and specific leaf area (SLA)) and below-ground traits (root average diameter (RD), root biomass (RB), specific root length (SRL), specific root area (SRA) and root tissue density (RTD)) in undisturbed and overgrazed meadows for all species, and separately for annuals and perennial species from 3000 to 3600m elevations.

PCA results revealed that undisturbed and overgrazed species were significantly separated along axis 1 at all elevations. PCA axis 1 had a strong positive loading for SRL, SRA, A and SLA for all common species at all elevations, but negative loading for RD, RB, RTD and H (Fig. 3 and Tables S2). PCA axis 2 was significantly positively associated with RB and RTD (Fig. 3, and Table S2). When performing PCA separately for annual and perennial species, we still found consistent PCA results for perennial species. However, we found different PCA results for annual species, because undisturbed and overgrazed species could not be significantly separated along axis 1 and axis 2 at all elevations (Fig. S1 and Table S2). Moreover loading for traits that was signficantly associated with PCA axis 1 and 2 were different among all elevations (Fig. S1 and Table S2).

**Fig. 3.**
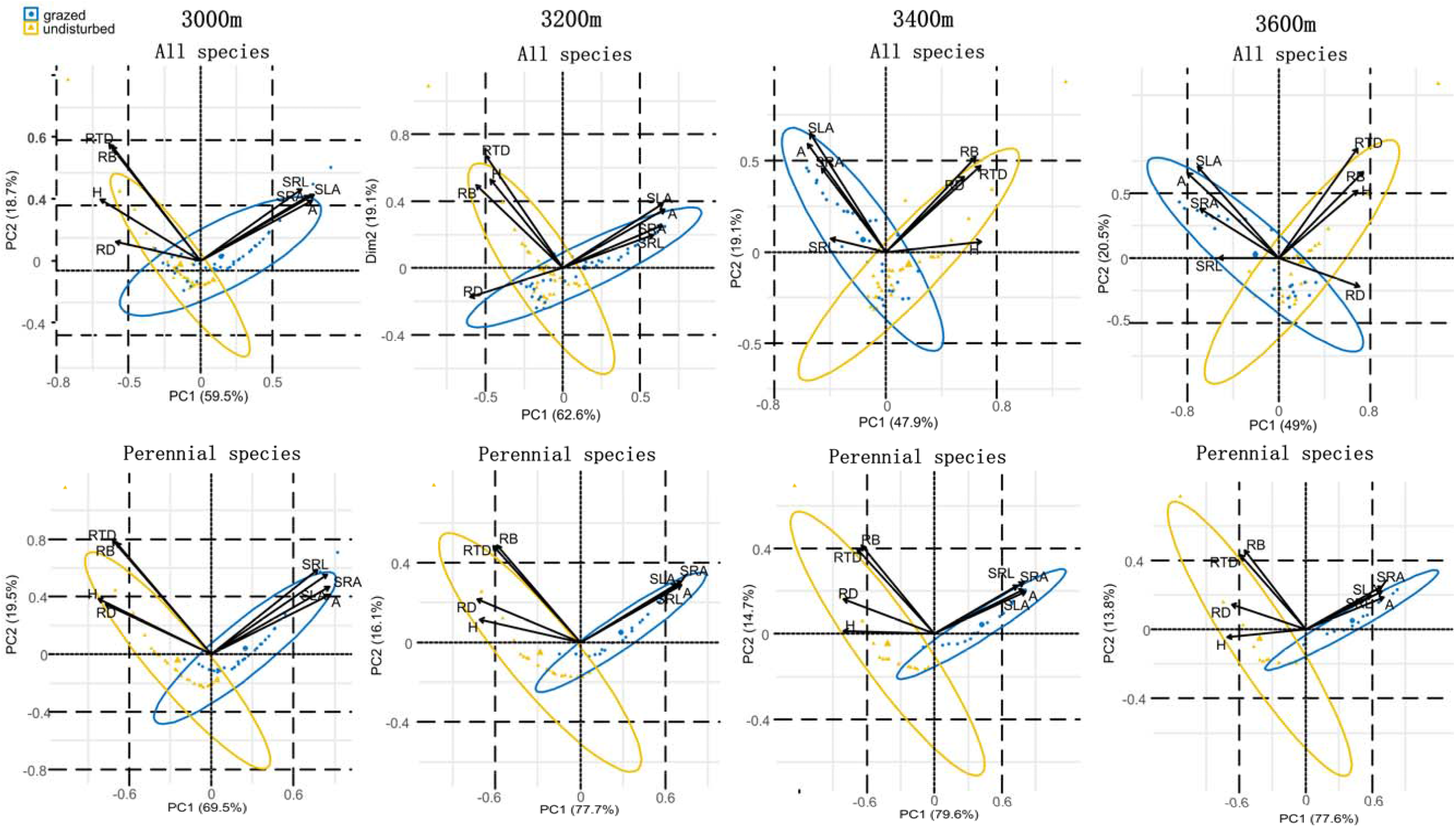
Principal component analysis for below-ground traits (root average diameter (RD), root biomass (RB), specific root length (SRL), specific root area (SRA) and root tissue density (RTD)) and above-ground traits (height (H), leaf maximum photosynthesis rate (A) and specific leaf area (SLA)) in undisturbed and overgrazed meadows for all species and perennial species from 3000 to 3600m elevations.

## Discussion

In this study, we quantified how overgrazing affect above-ground and below-ground traits; and whether the response of above- and below-ground traits to overgrazing varied with life form (annual versus perennial species). As we hypothesized, both above-ground and below-ground traits exhibited a resource acquisition *vs*. resource conservation trade-off for responding to overgrazing. The response of above- and below-ground traits to overgrazing differed between annual and perennial species, with perennial species determining the response of above- and below-ground traits to overgrazing.

### The response of above-ground and below-ground traits to overgrazing for all species at all elevations

A shift from resource conservation to resource acquisition at above-ground level have been widely found to tolerate grazing, which force species to have higher SLA, photosynthesis rate, but lower height after overgrazing than those in the undisturbed condition (Díaz *et al*., 2001; Spasojevic *et al*., 2010; Laliberté and Tylianakis, 2012; Li *et al*., 2017). It has also been found that this resource conservation vs. resource acquisition trade-off has also led to strong correlation among SLA, photosynthesis rate and height (Wright *et al*., 2004; Osnas *et al*., 2013). Here we observed consistent trait variations and correlation among SLA, photosynthesis rate and height for all species in both undisturbed and overgrazed meadows at all four elevations. This indicated that a trade-off between resource acquisition *vs*. resource conservation at above-ground level likely determines plant tolerance to overgrazing in Qinghai Tibetan plateau.

A number of studies have found that the trade-off between resource acquisition *vs*. resource conservation can also be found in below-ground traits, similar to what have been found in above-ground traits (Withington *et al*., 2006; Mommer and Weemstra, 2012; Reich, 2014; Roumet *et al*., 2016). For example, resource acquisition may need species to have high SRL, SRA, but low RTD, RB and RD to possess fast-growing, whereas, resource conservation may force species to have high RTD, RB and RD, but low SRL and SRA (Reich, 2014; Roumet *et al*., 2016). Likewise, this resource acquisition vs. resource conservation trade-off at below-ground level can also led to significant root correlations (Withington *et al*., 2006; Prieto *et al*., 2015; de la Riva *et al*., 2018). Indeed, we found consistent trait variations and correlations among below-ground traits (SRL, SRA, RTD, RD and RB) at all elevations. This, indicated below-ground traits have also a shift from resource conservation to resource acquisition to tolerate overgrazing as above-ground traits does.

In the undisturbed meadows, resource conservation can also force species to have low photosynthetic rates and SLA, which require conservative root structures (e.g., roots with high root tissue density, root diameter and biomass, and/or low specific root length and area) to ensure high overall conservation of resources (Eissenstat, 2002; Fort *et al*., 2013). In contrast, in the grazed meadow, resource acquisition tends to make species possess high photosynthetic rates and SLA, which requires roots with high specific root length and area, and low root tissue density, diameter and biomass to ensure sufficient nutrient supply to achieve fast plant growth (Reich, 2014; de la Riva *et al*., 2018). Thus, significant above-ground-below-ground traits relationship may be observed in both undisturbed and grazed meadows. Indeed, significant relationships between above- and below-ground traits have been observed in multiple studies in undisturbed meadows (Wardel *et al*., 2004; Liu *et al*., 2010; Valverde-Barrantes *et al*., 2017; de la Riva *et al*., 2018). However, nearly no study reported strong relationships between above-ground traits and below-ground traits after overgrazing. Here we observed consistently significant relationships between above- and below-ground traits in both undisturbed and overgrazed meadows at all elevations. Thus, above-ground traits might act as a proxy for below ground traits in revealing the response to overgrazing at the below-ground level.

Principal component analysis revealed that undistrbed and overgrazed species were significantly separated along axis 1, which was significantly positively associated with SRL, SRA, A and SLA, but negatively related RD, RB, RTD and H. Since high SRL, SRA, A and SLA represented high resource acquisiiton, whereas, high RD, RB, RTD and H indicated high resource conservation (Wright *et al*., 2004; Reich, 2014; de la Riva *et al*., 2018). Thus the shift from resource conservation to resource acquisition determine species distribution between undisturbed and overgrazed meadows.

### Different response of above-ground and below-ground traits to overgrazing between annual and perennial species

It has been universally found that grazing promotes annual over perennial species in grassland ecosystems (Díaz *et al*., 2007). Our study supports this trend where we found that after overgrazing perennial species decrease 2-2.3 times, whereas, annual species increase 1.2-1.9 times. Especially at 3600m elevation, after overgrazing, annual species is more than perennial species. Annuals tend to have traits associated with a higher resource acquisition strategy, while perennial species tend to have traits associated with a resource conservation strategy (Roumet *et al*., 2006; Zhang *et al*., 2018). As a result, the outcomes for trait variations, trait-trait correlations and PCA may vary between annual species and perennial species. Indeed, consistent trait variations, trait-trait correlations and PCA results for all species can merely be found in perennial species but not annual species at all elevations. Our data, on one hand indicated that the response of above- and below-ground traits to overgrazing indeed varied between annual and perennial species. On the other, this also indicated that species-specific responses to grazing are mainly determined by perennial plants (Dorrough *et al*., 2004).

Since species numbers for perennial species are nearly twice of those for annual species among common species between undisturbed and overgrazed meadow at 3000 and 3200m elevations, it is possible that consistent results for all species and perennial species, but not for annual species may be attributed to much more species numbers for perennial species from the perspective of statistic analysis. Thus species-specific responses to grazing may be not determined by perennial plants. However, at 3400m and 3600m elevations, although species numbers for annual species are more than those for perennial species among shared species between undisturbed and overgrazed madows, we still found consistent results for all species and perennial species, but not for annual species. Consequently, perennial species but not annual species determine the response of above- and below-ground traits to overgrazing.

As shown in Fig. 4, our data can be used to disentangle different responses to undisturbed and overgrazed condition between annual and perennial species. Due to N-limitation in undisturbed sites (Zhang *et al*., 2018), perennial species tend to have high resource conservation strategies. After overgrazing, perennial species are largely removed, whereas annual species largely increased after overgrazing. Thus overgrazing has positive effect on annual species, but negative effect on perennial species. This may force perennial species to change its intrinsic resource conservation into resource acquisition strategy to tolerate overgrazing. In contrast, annual species might be not necessary to make any response for overgrazing.

**Fig. 4.**
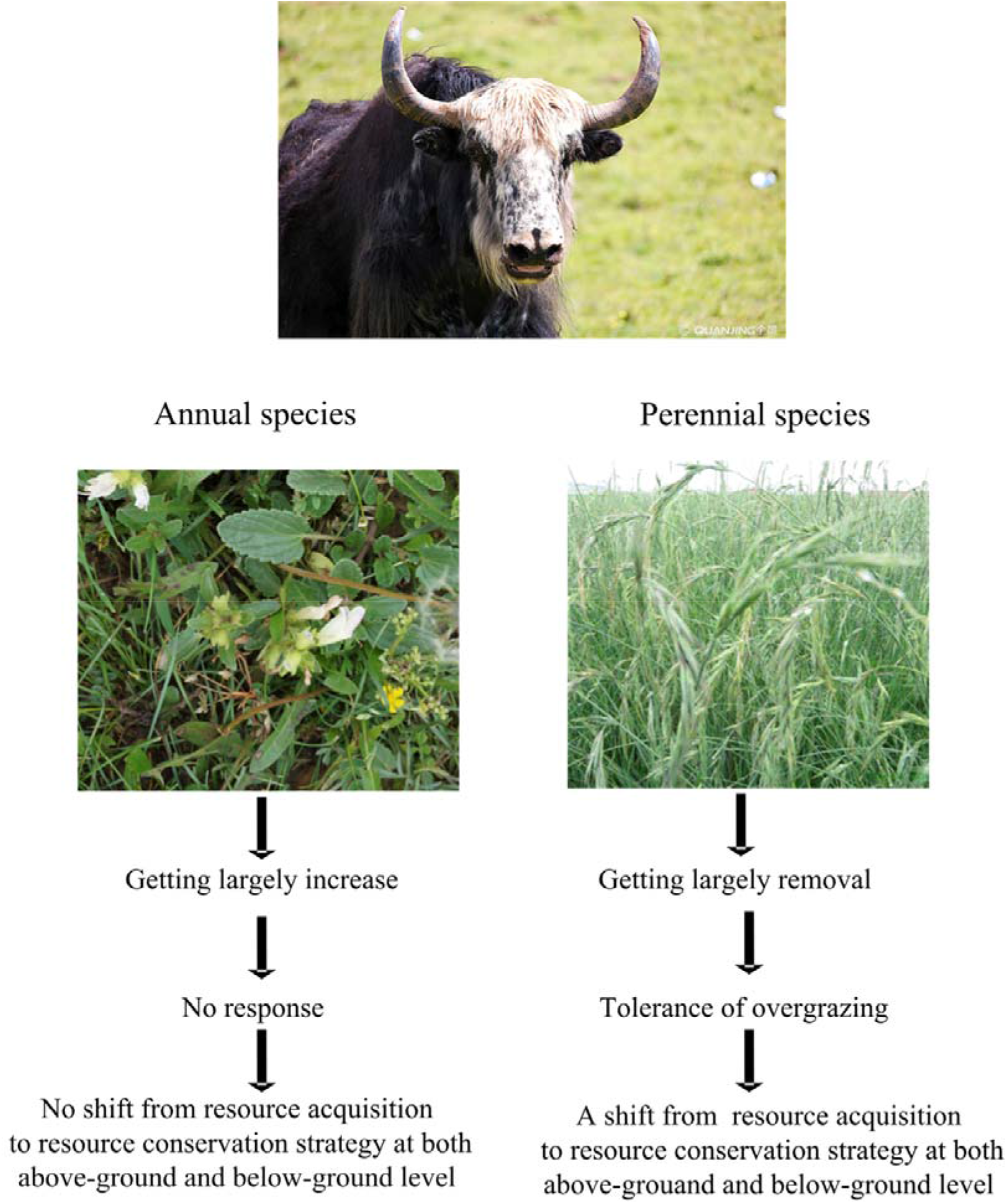
Hypothesized patterns of different responses of annual and perennial species to overgrazing.

## Conclusion

Together, our results suggested that both above-ground and below-ground traits exhibited similar resource conservation *vs*. resource acquisition trade-off for responding to overgrazing. Moreover, perennial species but not annual species determine the response of above- and below-ground traits to overgrazing. Our findings also have important implications for the management and recovery of overgrazed meadow ecosystems. First, since,as strong correlations among above-ground traits and below-ground traits can be found in overgrazed meadow, above-ground traits can be a good proxy for below-ground traits, and can be used to understand plant-soil feedbacks and the effects of below-ground traits on ecosystem functioning in overgrazed meadows. Second, since the shift from resource conservation to resource acquisition determine species distribution between undisturbed and overgrazed meadow, selecting species that have high resource acquisition can help the diversification of the overgrazed meadow. Likewise, selecting desirable species with high resource acquisition to undesirable weeds may help control weeds. Third, since perennial species determine the response of above-ground and below-ground traits to overgrazing, which in turn affect vegetation composition between undisturbed and overgrazed meadow, active seeding of perennial species may lead to strong competition between perennial species and annual species in the overgrazed meadow, which may in turn quickly restore overgrazed meadow to natural condition.

## Acknowledgments

This work was funded by the National Natural Science Foundation of China (31770469) and by a start-up fund from Hainan University (KYQD (ZR) 1876). The authors declare no conflict of interest.

